# Light exposure decreases infectivity of the *Daphnia* parasite *Pasteuria ramosa*

**DOI:** 10.1101/789628

**Authors:** Erin P. Overholt, Meghan A. Duffy, Matthew P. Meeks, Taylor H. Leach, Craig E. Williamson

**Affiliations:** Department of Biology, Miami University, Oxford, OH; Department of Ecology and Evolutionary Biology, University of Michigan, Ann Arbor, MI

**Keywords:** zooplankton, UV, endospore, pathogen

## Abstract

Climate change is altering light regimes in lakes, which should impact disease outbreaks, since sunlight can harm aquatic pathogens. However, some bacterial endospores are resistant to damage from light, even surviving exposure to UV-C. We examined the sensitivity of *Pasteuria ramosa* endospores, an aquatic parasite infecting *Daphnia* zooplankton, to biologically relevant wavelengths of light. Laboratory exposure to increasing intensities of UV-B, UV-A, and visible light significantly decreased *P. ramosa* infectivity, though there was no effect of spore exposure on parasitic castration of the host. *P. ramosa* is more sensitive than its *Daphnia* host to damage by longer wavelength UV-A and visible light; this may enable *Daphnia* to seek an optimal light environment in the water column where both UV-B damage and parasitism are minimal. Studies of pathogen light sensitivity help us uncover factors controlling epidemics in lakes, which is especially important given that water transparency is decreasing in many lakes.

## INTRODUCTION

Changing temperature and precipitation related to climate change are altering disease dynamics. One factor that plays a role is declining water transparency, since ultraviolet light penetration into lakes has germicidal effects (Williamson et al. 2017). Thus, by decreasing light penetration in lakes, climate change has the potential to promote epidemics.

However, while we know that many microbes are harmed by exposure to light, we also know some tolerate light remarkably well. Endospores, a resting stage found only in Gram positive bacteria of the group Firmicutes, are highly resistant to disinfecting techniques (Nicholson *et al.*, 2000), including high levels of UV-C radiation (Newcombe *et al.*, 2005). Surprisingly though, in the endospore form, the highly studied *Bacillus* showed decreased survival across a wide range of exposures, from UV-B to full sunlight (Xue and Nicholson, 1996). Other pathogens have shown a similar sensitivity to longer wavelengths of light. For example, an aquatic pathogen - the fungus *Metschnikowia* -was sensitive to solar radiation even in the absence of UV, and field surveys showed larger epidemics in less transparent lakes (Overholt *et al.*, 2012).

Light can harm pathogens, and climate change is altering light regimes in lakes. Thus, investigations into how aquatic pathogens and their hosts respond to light are needed to better understand and predict disease dynamics. Here we test whether endospores of the virulent bacterial pathogen *Pasteuria ramosa* are sensitive to biologically relevant wavelengths of light, and if light exposure decreases its ability to lower fecundity in infected *Daphnia* hosts.

## METHODS

We exposed *P. ramosa* to different environmentally realistic light conditions in the laboratory and measured subsequent pathogen infectivity and host reproduction in *Daphnia dentifera*. Shallow quartz dishes containing 25 mL aliquots of *P. ramosa* spores (2000 spores mL^−1^) were placed on a rotating wheel (2 rpm) for 12 h at 24°C in a UV-lamp phototron and exposed to different levels of biologically relevant UV-B, and photorepair radiation (PRR, comprised of UV-A and visible light), which stimulates repair of UV-damaged DNA (Williamson *et al.*, 2001). In experiment 1, we examined the infectivity of *P. ramosa* under 10 intensities of PRR (8 replicates per treatment). Experiment 2 used a two-way factorial design to measure the effects of light wavelength and intensity on pathogen infectivity and host fecundity. Spores were exposed to either UVB and PRR or visible light only at nine intensity levels. For the exposure, spores (2000 spores mL^−1^) were aliquoted into five replicates of each treatment (intensity x light type). (See supplement for additional details.)

Following exposure in the phototron, dishes were removed and a single, week-old *D. dentifera* neonate was placed in each dish with the exposed *P. ramosa* spores. After three days, *D. dentifera* were transferred to 50 mL beakers and filled with 30 mL of spore-less, filtered lake water. In both experiments, offspring were removed during water changes. In experiment 2, neonates were quantified during water changes and eggs in the brood chamber counted on day 25. After 25 days, individuals were examined for infection.

We used a binomial logistic regression model to test the effects of increasing light intensity on infectivity for each light treatment. A two-way ANOVA was used to test the effect of light treatment and intensity on the number of neonates produced. Analyses were conducted using R version 3.4.4 (R Core Development Team).

## RESULTS

Light exposure greatly decreased parasite infectivity: In both experiments, the highest rates of infection were in the dark (0% light) and decreased with increasing light exposure of the pathogen (Fig. 1). The fecundity of *Daphnia* also increased with increasing exposure of *P. ramosa* to light, but, for hosts that became infected, there was no difference in the number of neonates produced (Fig. 2).

**Fig. 1.**
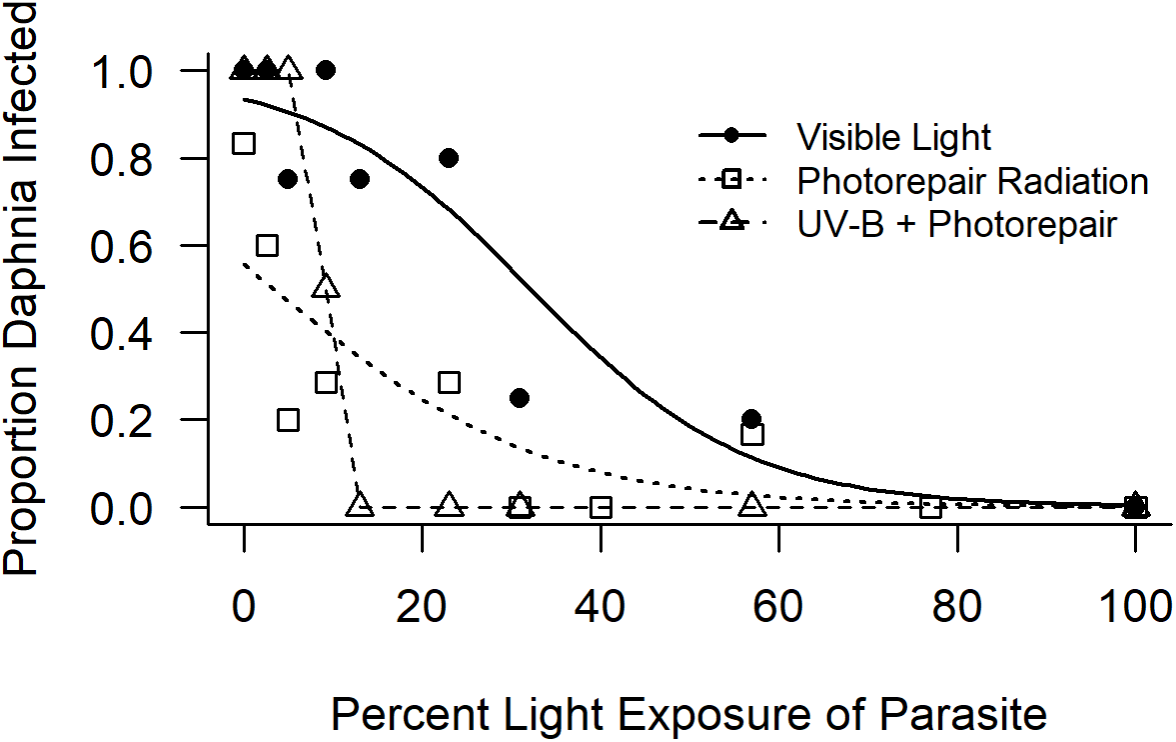
Proportion of *Daphnia* infected at each exposure level of the pathogen to visible light, photorepair radiation, or UV-B + photorepair. Lines represent the fitted logistic regression model for the visible light and photorepair radiation treatments. When UV was blocked, the proportion of *D. dentifera* infected significantly decreased with increasing exposure of the pathogen (experiment 2 Visible Light, p < 0.01); the logistic model fit indicated that there was an 8.0% decreased probability of infection for every 1% increase in light exposure. We also found a significant negative relationship between light intensity and infectivity when the pathogen was exposed to increasing levels of photorepair radiation (experiment 1 Photorepair Radiation, p < 0.01); the logistic model indicated that for every 1% increase in PRR exposure the probability of *D. dentifera* infection decreased by 6.5%. When exposed to the full spectrum of light (experiment 2 UV-B + Photorepair), there was also a decline in infectivity; this model would not converge due to sharp cut off in proportion infected, so the line does not represent the fitted model, but instead simply connects the points.

**Fig. 2:**
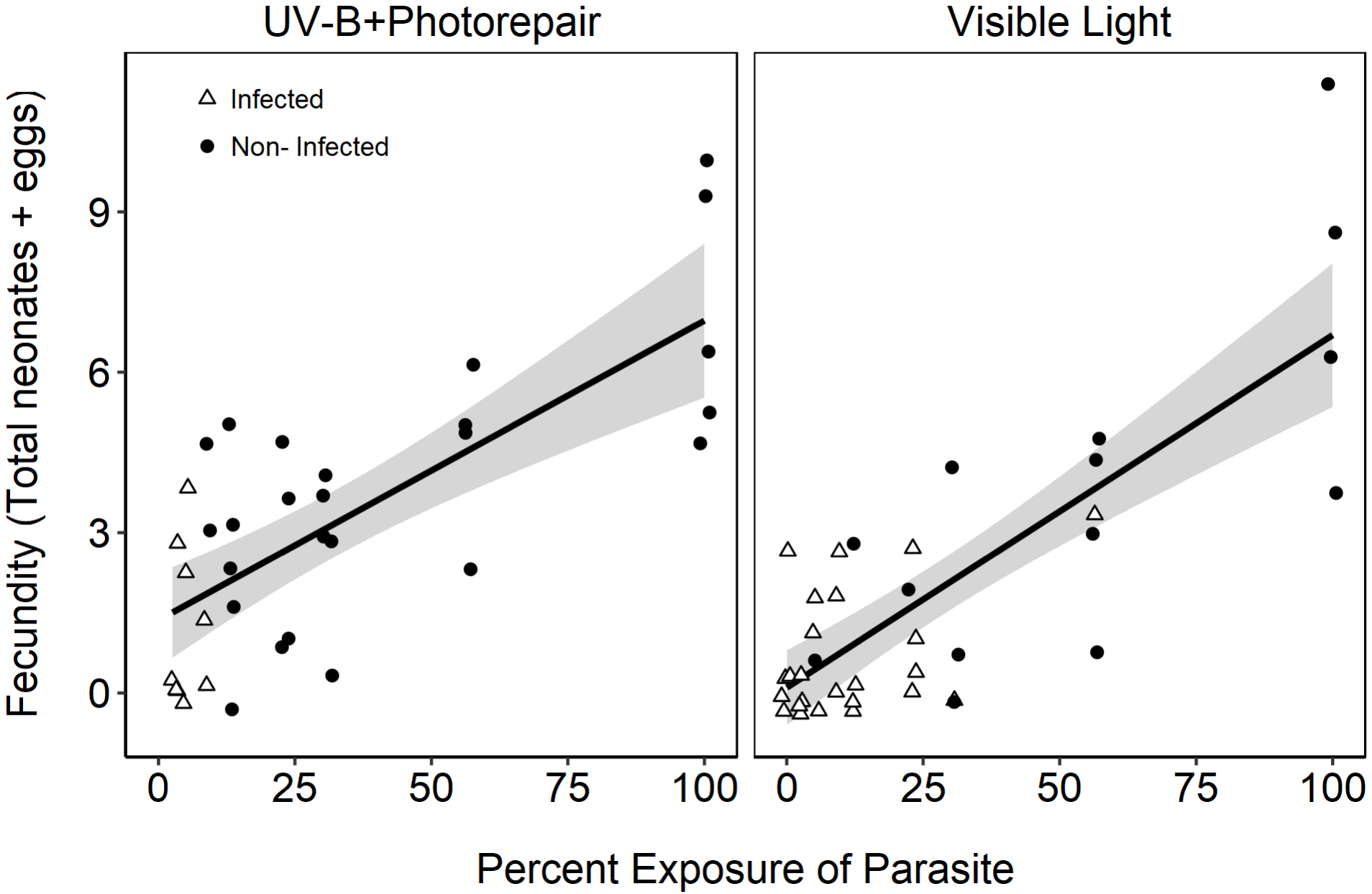
Fecundity (total neonates plus eggs on day 25) of infected (triangles) and uninfected (circles) *Daphnia* in experiment 2. Overall, *Daphnia* fecundity increased when spores were exposed to higher intensities (intensity p < 0.001, light treatment p = 0.03), but, for hosts that became infected, there was no difference in the number of neonates produced in the different treatments (intensity p = 0.37; light treatment p= 0.57). The lines represent linear regressions with shared 95% confidence intervals for infected and uninfected individuals.

## DISCUSSION

We found that endospores of *P. ramosa* are surprisingly susceptible to longer wavelength UV-B, UV-A and even visible light. This influence of light on spores benefitted *Daphnia* via impacts on infection: hosts that were exposed to spores that had been exposed to more light were less likely to become infected and, therefore, produced more offspring. However, when just infected hosts were considered, spore light exposure did not alter host reproduction.

Other aquatic pathogens are sensitive to longer wavelengths of light as well. For example, *Cryptosporidium* cysts lost infectivity following exposure to UV (Connelly *et al.*, 2007; King *et al.*, 2008) and visible light (Connelly *et al.*, 2007). Natural sunlight caused additional sublethal effects on the protein secretion required by *Cryptosporidium* for attachment to its host (King *et al.*, 2010). The fungal parasite *Metschnikowia*, which can be found in the same lake systems as *P. ramosa*, was also sensitive to both short wavelength UV-B as well as longer wavelength UV-A and visible light in both laboratory and field studies (Overholt *et al.*, 2012). Another field study suggested that *Pasteuria* was susceptible to solar radiation; however sensitivity to visible light was not specifically tested (C. L. Shaw, unpublished data).

In our study, *P. ramosa* exhibited decreased infectivity even under long wavelength UV-A and visible light, in the absence of UV-B, though shorter wavelengths overall caused the greatest decrease in infectivity. In contrast, these longer wavelengths benefit the host, *Daphnia*, by stimulating photorepair of DNA damage (Sancar, 1994). Since the same wavelengths that damage the pathogen can benefit the host, *Daphnia* may be able to find a refuge from disease and damaging UV-B at intermediate depths in the water column where UV-A and visible light levels are high, but damaging UV-B is less intense.

## CONCLUSIONS

Lakes in many regions are experiencing lower intensity light regimes due to increased dissolved organic matter inputs and/or eutrophication (Monteith *et al.*, 2007; Solomon *et al.*, 2015; Strock *et al.*, 2017; Williamson *et al.*, 2017; Williamson *et al.*, 2015). Our finding that the common bacterial pathogen *P. ramosa* is sensitive to both UV and visible light suggests that decreases in lake transparency through “browning” and/or “greening,” may allow for increased *P. ramosa* prevalence.

## SUPPLEMENTARY DATA

Supplementary data will be available online after publication.

## ACKNOWLEDGMENTS

We thank Rebecca Healy for preliminary *Pasteuria* work.

## SUPPLEMENTAL METHODS

*D. dentifera* were isolated from Bishop Lake (Livingston County, MI, USA) and maintained on a diet of *Ankistrodesmus falcatus* at 24°C on a 16:8 light: dark cycle throughout the experiment. A single clone of the host species was used to standardize the susceptibility of the host organism to the bacterial *P.ramosa* strains.

PRR was supplied by two 40-W UV-A bulbs (Q-Lab QUV UV-A, Q-Lab, Cleveland, OH) and two fluorescent, 40 W, cool white tube bulbs. Light intensity was manipulated using mesh screens as neutral density filters. A total of ten intensity treatments, with eight replicates each, were used in experiment 1 (0%, 2.6%, 5%, 9.2%, 23%, 31%, 40%, 57%, 77%, 100%).

In experiment 2, PRR and UV-B were crossed factorially. Covering the UV-B lamp (Spectronics XX15B, Rochester, New York) with acetate removed wavelengths less than 295 nm, allowing only biologically relevant wavelengths to be transmitted. The addition of light filters on individual dishes either transmitted UV (creating the “UV-B + photorepair” treatment) or blocked all UV (creating the “visible light” treatment). In the UV-B + photorepair treatment, a plastic filter (printed acetate, transmitting 91% photosynthetically active radiation (PAR) 400–700 nm, 87% UV-A 320–399 nm, and 70% UV-B 295–319 nm, in air) allowed full-spectrum exposure while keeping the starting intensity between UV-B + photorepair and visible light treatments similar. In the visible light treatment, Courtgard (CP Films, Inc., Martinsville, VA, USA) transmitted visible light, but blocked most UV (in air, transmits 87% of 400–700 nm PAR, 0% UV-B 295–319 nm and 6% of UV-A 320– 399 nm). Similar to experiment 1, mesh screens acting as neutral density filters were again used to create nine intensity levels (0%, 2.6%, 5%, 9.2%, 13%, 23%, 31%, 57%, 100%).

When *D. dentifera* were exposed to spores, all dishes were placed in an incubator at 24°C on a 16:8 light (cool white bulbs): dark cycle. Individual *D. dentifera* were fed *A. falcatus* (1 x 103 cells mL-1) daily for three days. After three days, individuals were moved to clean, spore-free water. Individuals were then fed daily, and the water was changed every three days.

